# Cyanogenic glucoside production in cassava: The comparable influences of varieties, soil moisture content and nutrient supply

**DOI:** 10.1101/649236

**Authors:** Matema L.E. Imakumbili, Ernest Semu, Johnson M.R. Semoka, Adebayo Abass, Geoffrey Mkamilo

## Abstract

Varieties and soil moisture content are the two agronomic factors mostly pointed out as influencers of cyanogenic glucoside production in cassava. The role of soil nutrient supply is however often overlooked or minimised, despite its known influence on cyanogenic glucoside production. A pot experiment was hence carried out to determine whether soil nutrient supply had an equal influence on cyanogenic glucoside production in cassava, as varieties and soil moisture content. The cassava varieties, *Kiroba* (a sweet cassava variety) and *Salanga* (a bitter cassava variety), were used in the experiment, together with three soil moisture treatments that respectively induced severe moisture stress, moderate moisture stress and no moisture stress (optimal soil moisture conditions where plants were kept well-watered). The soil nutrient treatments used depicted conditions of low (no fertiliser), moderate (25 N mg, 5 P mg, 25 K mg /kg) and high (25 N mg, 5 P mg, 25 K mg /kg) nutrient supply. A sole K treatment was also included (25 K mg/kg). Total hydrogen cyanide (HCN) levels in cassava leaves were used to indicate the effects of the three factors on cyanogenic glucoside production. The results of the study showed that nutrient supply had a significantly (p < 0.001) equal influence on cyanogenic glucoside production, as varieties (p < 0.001) and soil moisture content (p < 0.001). Cyanogenic glucoside production was however found to be differently influenced by soil moisture content (M) and nutrient supply (N) in both *Salanga* (M×N, p = 0.002) and *Kiroba* (M×N, p < 0.001). Leaf HCN levels of unfertilised *Salanga* and *Kiroba* were respectively increased by 1.8 times and 2.7 times their levels under optimal soil moisture conditions. Thus, under severe moisture stress, low soil fertility was found to have an increasing effect on leaf HCN levels in both varieties. A high supply of N, P and K, however also had an increasing effect on leaf HCN in both varieties regardless of soil moisture conditions. Leaf HCN levels in *Salanga* ranged from 95.5 mg/kg to 334.5 mg/kg and in *Kiroba* they ranged from 39.3 mg/kg to 161.5 mg/kg, on a fresh weight basis. The study managed to demonstrate that soil fertility had an equally important influence on cyanogenic glucoside production, just like varieties and soil moisture content. The study also showed that the effects of nutrient supply on cyanogenic glucoside production in various cassava varieties is dependent on changes in soil moisture content and vice versa.

## Introduction

Although it is well-known that soil nutrient supply influences cyanogenic glucoside production in cassava (*Manihot esculenta* Crantz) [1–4], the importance of its role is not given as much attention. For example, despite the known low soil fertility of the predominantly sandy soils in areas affected by cassava cyanide intoxication in sub-Saharan Africa [5–7], more attention is given to the agronomic effects of moisture stress and to the bitter cassava varieties commonly planted by farmers living in these areas of low agricultural potential [5,8–11]. Like in this example, soil moisture content and variety type are usually considered as the main influencers of cyanogenic glucoside production in cassava, while the role of soil nutrient supply (soil fertility) is often overlooked or is given less importance.

In one study, reductions in cassava root cyanogenic glucoside levels were separately achieved by both mulching and fertiliser application [12], suggesting the equally important influence of nutrient supply on cyanogenic glucoside production in comparison to soil moisture content. Reductions in cyanogenic glucosides were however less pronounced when cassava was both mulched and fertilised [12], revealing a slight cancellation of the individual effects of nutrient supply and soil moisture content on cyanogenic glucoside production when their effects were combined. The importance of the less considered role of soil nutrient supply was hence once again highlighted in the odd result of the combined effects of soil moisture content and nutrient supply. The cancelling of the individual effects of the two environmental factors (soil moisture content and nutrient supply) could explain why cyanogenic glucoside production is not always influenced by soil nutrient supply [13]. The different effects that agro-climatic environments have on cyanogenic glucoside production in cassava varieties [14–16] could also be a result of the little understood effects of soil nutrient supply and its interaction with other agronomic factors.

In this study we tried to shed light on the influence that soil nutrient supply has on cyanogenic glucoside production in cassava, in comparison to the influences of varieties and soil moisture content. The hypothesis tested was that ‘*soil nutrient supply has no influence on cyanogenic glucoside production, unlike variety and soil moisture content*’. A pot experiment where all three factors were included was used to investigate their comparable influence on cyanogenic glucoside production. The cyanogenic glucoside content of cassava leaves was used to determine the effects of varieties, soil moisture content and nutrient supply on cyanogenic glucoside production. The cyanogenic glucoside content of cassava leaves was used given that leaves are the main sites of cyanogenic glucoside synthesis in cassava. Cyanogenic glucoside levels in cassava leaves would thus be a good indicator of the effects of the three factors on cyanogenic glucoside production in cassava plants. This off course does not ignore the fact that cyanogenic glucosides accumulate differently in various parts of cassava plants, but helps to show a general trend on the expected effects on cyanogenic glucoside production in the entire plant.

## Materials and methods

### Location, experimental design and treatments

The pot experiment was a 2×3×4 factorial combination of two cassava varieties, three soil moisture levels and four different nutrient supply treatments. The experiment was laid out using the Randomised Complete Block Design (RCBD) and was replicated five times. The experiment was replicated five times because a minimum of four plants is required when determining the cyanogenic glucoside content of leaves of a particular cassava variety growing in the same field or under a similar set of treatments like in this case [17]. Only four replicates were however considered as one block was removed due to incorrect labelling of treatments. Blocking was necessary due to the differential lighting in the screen house. Each block consisted of 24 treatments (pots). The experiment was carried out for 90 days from 24^th^ October 2015 to 25^th^ January 2016, at Sokoine University of Agriculture (SUA) (S 6°51’13”, E 37°39’26”) in Morogoro district in Tanzania.

### Cassava varieties

A sweet cassava variety and a bitter cassava variety were used in the pot experiment. The sweet variety used was an improved cassava variety called *Kiroba*. This variety is grown by a number of farmers in Mtwara region of Tanzania, because of its high yielding ability and also for its ability to resist diseases and to tolerate pests. The bitter variety used was a local cassava variety called *Salanga. Salanga* is commonly associated with cyanide intoxication in konzo-affected areas of Mtwara region. *Kiroba* was collected from Naliendele Agricultural Research Institute (NARI) (S 10°21’22”, E 40°09’59”), while *Salanga* was collected from Kitangari village (S 10°39’01”, E 39°20’01”) in Newala district in Tanzania. All cassava stem cuttings were collected from mature, ready to harvest and visibly healthy plants.

Previously rooted cassava plantlets were used to establish the pot experiment. To produce the rooted plantlets the mature stem cuttings of the two collected cassava varieties were first grown in a nursery using rapid cassava multiplication methods [https://dx.doi.org/10.17504/protocols.io.z9cf92w] [18]. Short stem cuttings of about 10 cm long for each collected cassava variety were densely planted at a spacing of 10 cm x 10 cm in beds with soil of low fertility and with no fertiliser application history. The cuttings were allowed to sprout and left growing until their shoots were 15 cm long. The shoots were then cut-off and rooted in distilled water to produce the rooted plantlets. It took about one month for the shoots to root in distilled water.

### Soil nutrient treatments

The role of nutrient supply was investigated using a varied supply of the three nutrients that are often deficient in most African soils, that is, nitrogen (N), phosphorous (P) and potassium (K). The nutrient treatments were formulated based on applications of low, moderate and high levels of N, P or K, which allowed the investigation of the effects of improved N, P and K supply on cyanogenic glucoside production in cassava. The first soil nutrient treatment was a control, in which no nutrients were applied. The second nutrient treatment was a K only nutrient treatment, in which K was solely applied at the rate of 25 mg/kg. A sole K nutrient treatment was included, given the reported reducing effects of K fertiliser on cassava cyanogenic glucosides [19]. The third nutrient treatment was a moderate NPK treatment, with N, P and K applied at rates of 25 mg/kg, 5 mg/kg and 25 mg/kg, respectively. The fourth nutrient treatment was a high NPK treatment, with N, P and K supplied at rates of 50 mg/kg, 13 mg/kg and 50 mg/kg, respectively. The NPK rates used to formulate the nutrient treatments were based on common field based NPK fertiliser rates applied to cassava. Although there are slight differences, the field based NPK fertiliser rates were based on the general recommended NPK fertiliser rate for cassava production (100 N kg, 22 P kg 83 K kg /ha or 100 N kg, 50 P_2_O_5_ kg 100 K_2_O kg /ha) [20]. The pot based NPK fertiliser rates expressed on a per kg soil basis, are given below, together with their equivalent field based NPK rates (Table 1).

**Table 1.**
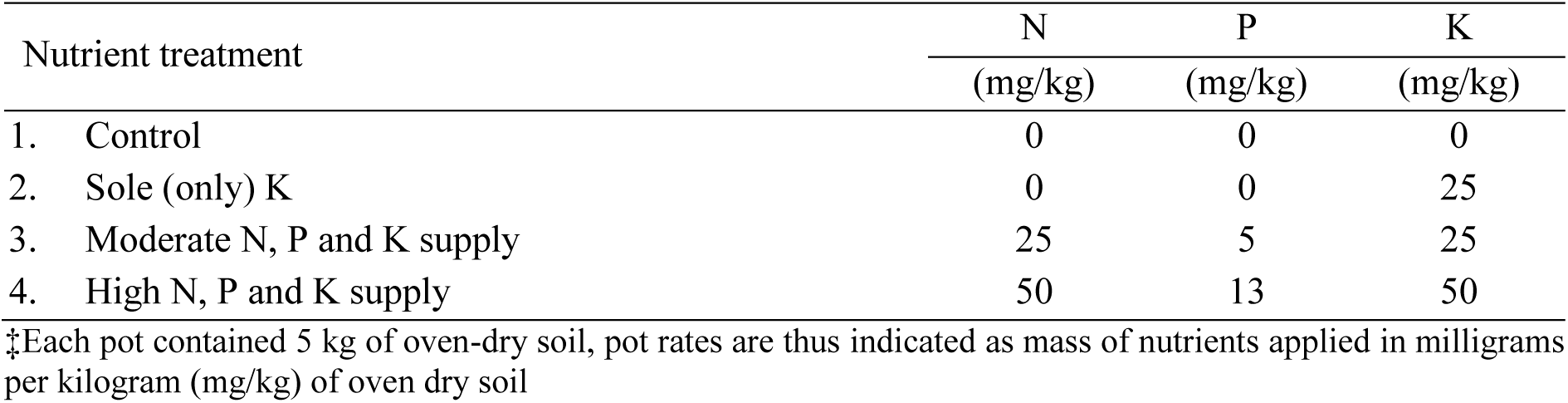
Nutrient treatments used in the pot experiment.

| Nutrient treatment | N | P | K |
| --- | --- | --- | --- |
|  | (mg/kg) | (mg/kg) | (mg/kg) |
| 1. Control | 0 | 0 | 0 |
| 2. Sole (only) K | 0 | 0 | 25 |
| 3. Moderate N, P and K supply | 25 | 5 | 25 |
| 4. High N, P and K supply | 50 | 13 | 50 |

‡Each pot contained 5 kg of oven-dry soil, pot rates are thus indicated as mass of nutrients applied in milligrams per kilogram (mg/kg) of oven dry soil

The fertilisers; urea (CO(NH_2_)_2_), triple super phosphate (TSP) (Ca(H_2_PO_4_)_2_.H_2_O) and muriate of potash (MOP) (KCl), were used to supply N, P and K. All the KCl and TSP were mixed into the soil before planting [https://dx.doi.org/10.17504/protocols.io.4engtde], while the urea was applied in solution form [21], in two split applications at two and six weeks after planting [https://dx.doi.org/10.17504/protocols.io.4ifgubn].

### Soil moisture treatments

All pots were kept well-watered (maintained at field capacity) during the first 69 days after planting (DAP). The soil moisture treatments were then only begun at 70 DAP and were maintained for the next 20 days, that is, until 90 DAP. The selected soil moisture treatments were intended to keep some plants well-watered, others moderately stressed and the rest severely stressed; this was achieved by irrigations which kept the soil in the treatment pots at 100%, 60% and 30% field capacity (FC), respectively [22–24]. The amount of water needed to bring the soils in the pots to complete field capacity was determined by merging two methods [https://dx.doi.org/10.17504/protocols.io.2xdgfi6] [22,23,25].

At 70 DAP, pots under the 100% FC moisture treatment were brought to field capacity each day, while pots under the 30% FC and 60% FC moisture levels were allowed to lose water until they were slightly below their needed respective field moisture levels. Once each pot had attained a slightly lower moisture content than 30% FC or 60% FC it was re-watered and maintained at the respective moisture level by daily replacement of the lost water for 20 days. Daily weighing of pots to determine the amount of water lost in each pot before re-watering was done around 07:00 hours each morning. The amount of moisture replaced in each pot took into consideration the weights of the pot, soil and plant [22].

### Soil collection and preparation for potting

To properly investigate the effects of nutrient supply on cyanogenic glucoside production, a nutrient poor soil was used in the pot experiment. Soil of this nature was collected from Soga village (S 6°49’54”, E 38°51’49”), located in Kibaha district in Tanzania. Soils in Kibaha district are predominantly Ferralic Cambisols and have an inherently low soil fertility [26,27]. Ten different points were selected from a field prior to soil collection. After removing surface litter, top soil was collected from the 10 selected points from a depth of 0 - 20 cm [https://dx.doi.org/10.17504/protocols.io.2eygbfw] [21,28]. The soil collected was packed in polypropylene sacks and transported to SUA where the experiment was to be conducted.

Before setting up the experiment, the collected soil was thoroughly mixed together and passed through an 8 mm sieve to facilitate drying and also to further remove stones, clods and trash. A 5 mm sieve should have however been used for this purpose. The soil was then spread and left to air-dry in an enclosed, dust free room for more than two weeks. The soil was turned from time to time to facilitate quicker and more even drying. When sufficiently dry the moisture content of the soil was determined gravimetrically [29]. The gravimetric moisture content of the soil was then used to calculate the amount of air-dry soil that would have to be weighed to give 5 kg of oven-dry soil to be placed in each pot. Uniform plastic pots with a 5 L capacity were used in the experiment. The soil was placed in the pots one day before planting. The soil was watered to field capacity on this day and left to thoroughly saturate with water overnight. Planting was carried out early in the morning on the following day. Each pot had a saucer below it to collect any excess water that drained from it. This however hardly happened. Capturing excess water on saucers, helped avoid nutrient losses from leached nutrients. The pots had holes in the bottom to let out excess irrigation water. The water collected on the saucers was placed back in the pot, preventing nutrient losses.

### Soil chemical and physical characteristics of the potting soil

The soil was analysed for organic carbon (OC), soil reaction (pH), total N, available P, available K, exchangeable calcium (Ca), exchangeable magnesium (Mg), available sulphur (S), extractable zinc (Zn), copper (Cu) and iron (Fe) and for soil texture before the experiment was set-up [https://dx.doi.org/10.17504/protocols.io.x72frqe]. All soil analysis procedures were carried out as follows [29]: pH in H_2_O, using a 1:1 soil to water ratio; OC using the Walkley and Black method; N was determined by micro-Kjeldahl digestion; P using the Bray No. 1 method; Sulphate-S using calcium phosphate [Ca(H_2_PO_4_)_2_] extracting solution; K, Ca and Mg in 1N ammonium acetate (NH_4_OAc) buffered at pH 7; extractable Zn, Cu and Fe using diethylenetriaminepentaacetic acid (DTPA); and texture by the hydrometer method. The results of the soil analysis are shown in Table 2, together with the respective ratings for their suitability for cassava production. The soil was a loamy sand with 86% sand, 11% clay and 2% silt [30].

**Table 2.** Soil chemical properties of the potting soil.

| Parameter | Value | Rating‡ | Sufficiency range or critical level | Reference |
| --- | --- | --- | --- | --- |
| pH | 5.80 | m | 4.5 – 7.0 | [18] |
| OC (%) | 0.35 | vl | 4.0 – 10.0 | [31] |
| N (%) | 0.06 | vl | 0.20 – 0.50 | [31] |
| P (mg/kg) | 3.54 | l | < 4.2 | [32] |
| K (cmol/kg) | 0.14 | l | 0.15 – 0.25 | [18] |
| Ca (cmol/kg) | 3.04 | m | 1.0 – 5.0 | [18] |
| Mg (cmol/kg) | 0.08 | vl | 0.40 – 1.00 | [18] |
| S (mg/kg) | 1.27 | l | < 6.0 | [31] |
| Zn (mg/kg) | 0.82 | l | 1.0 – 3.0 | [33] |
| Cu (mg/kg) | 0.70 | m | 0.3 – 0.8 | [33] |
| Fe (mg/kg) | 25.12 | vh | 4.0 – 6.0 | [33] |

‡vl, l, m and vh stand for very low, low, medium and very high

## Cultural practices

### Correction of nutrient deficiencies and pest control

The soil was deficient in N, P and K which was good for getting clear responses to N, P and K treatments (Table 2). The soils were however also deficient in Mg, S and Zn which had to be corrected, in order to obtain clear effects with N, P and K supply. Deficiencies in Mg and S were corrected using magnesium sulphate (MgSO_4_.7H_2_O), applied at a rate of 50 kg Mg/ha [20] by mixing 25 mg Mg/kg (simultaneously adding 32.5 mg S/kg) to the soil of each pot before planting. This was done at the same time that the TSP and MOP were also being added. A 2% solution of YaraVita Zintrac (700 g Zn/L; as ZnO), a chelated foliar fertiliser, was used to correct the Zn deficiency. It was applied to cassava at one month after planting and again at two months after planting. To keep insect pests away, the broad spectrum insecticide, Dursban (C_9_H_11_C_l3_NO_3_PS) was used. Dursban was always mixed with the ZnO foliar solution and was thus sprayed (applied) at the same time as the ZnO foliar solution.

### Irrigation water

Tap water was used to irrigate the plants throughout the entire experiment. The electrical conductivity of the water (EC_w_) was 0.007 dS/m and it had a pH of 6.58, on average. The water had a nitrate-nitrogen (NO_3_–N) content of 5.60 mg/L and only traces of phosphate-phosphorous (PO_4_–P). It also on average had 0.01, 0.16, 0.25 and 0.03 meq/L of K, Na, Ca and Mg, respectively. All parameters measured were within permissible levels for irrigation water [34,35]. In particular, there were negligible levels of N, P and K in the water making their additional contribution to nutrient supply very minimal.

### Determination of total hydrogen cyanide in cassava leaves

Leaf sampling for cyanide analyses was carried out at 91 DAP, which was the day that the experiment was also ended. Sample collection was carried out early in the morning around 07:00 hours, while ambient temperatures were low. The first fully-expanded leaf from the top of each cassava plant plus two leaves below it were picked during sample collection [https://dx.doi.org/10.17504/protocols.io.2dbga2n] [17]. The picrate paper method was used to determine the total HCN content of cassava leaves [https://dx.doi.org/10.17504/protocols.io.2dxga7n] [36,37]. After sample incubation, the reacted picrate papers were each later separately eluted in 5 ml of water and the absorbance of the picrate solution produced measured at the 510 nm wavelength using a spectrophotometer. The determined total HCN levels in the cassava leaves were expressed in mg/kg, on a fresh weight basis using Equation 1.

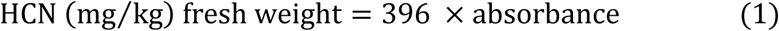

### Statistical analysis

The data collected was first analysed using a three-way analysis of variance (ANOVA) [38] to generally compare the individual effects of varieties, soil moisture content and nutrient supply on cyanogenic glucoside production in cassava. The data was then split by variety type and analysed using a two-way ANOVA, to determine the effects of soil moisture content and nutrient supply on cyanogenic glucoside production in each variety. Mean separation was carried out using the Tukey’s mean separation test at the 5% probability level. The data was screened for the identification of outliers which were removed before any analysis was carried out. All statistical analysis were carried out using GenStat Edition 14.

## Results and discussion

### Comparing the effects of varieties, soil moisture content and nutrient supply on leaf cyanogenic glucoside production

The F-test probability values obtained for the three-way ANOVA on the effects of varieties, soil moisture content and nutrient supply on cyanogenic glucoside production in cassava leaves are given in Table 3, where it can be seen that varieties, soil moisture content and nutrient supply had all significantly influenced cyanogenic glucoside production in cassava leaves. This showed that each factor had an important influence on cyanogenic glucoside production in cassava, confirming that soil nutrient supply has an equally important role as varieties and soil moisture content on cyanogenic glucoside production. The significant interaction effects of varieties, soil moisture content and nutrient supply (V×M×N, p < 0.05) (Table 3), additionally showed that no single factor (varieties or soil moisture content or nutrient supply) was responsible for the observed changes in leaf cyanogenic glucoside content. The effects of each factor on leaf cyanogenic glucoside content could thus only be explained by what had happened to the other two factors.

**Table 3.** F-test probability values for the three-way ANOVA on the effects of varieties, soil moisture content and nutrient supply on cassava leaf HCN content.

| Factor | p-value |
| --- | --- |
| Varieties (V) | < 0.001 *** |
| Soil moisture content (M) | < 0.001 *** |
| Nutrient supply (N) | < 0.001 *** |
| V×M | 0.398 NS |
| V×N | < 0.001 *** |
| M×N | 0.071 NS |
| V×M×N | < 0.001 *** |

\*\*\* Significant at $p < 0.001$ , \*\* significant at $p < 0.01$ , \* significant at $p < 0.05$ and NS not significant ( $p > 0.05$ ).

### Effects of soil moisture content and nutrient supply as observed for each cassava variety

The data was split by variety type to help better understand the significant V×M×N interaction effect obtained from the three-way ANOVA. Separate two-way ANOVA were hence carried out to determine how leaf cyanogenic glucoside production had been influenced by soil moisture content and nutrient supply in both *Salanga* and *Kiroba*. The F-test probability values obtained for the two-way ANOVA’s are shown in Table 4, where it can be seen that soil moisture content and nutrient supply had both significantly influenced leaf cyanogenic glucoside contents in the two varieties. The equally important influence of soil nutrient supply was once again shown and this time it is clear that its role is important in more than one cassava variety. Both varieties also had a significant M×N interaction (p < 0.05), which revealed that the effects of soil moisture content on leaf cyanogenic glucoside production in each variety were dependent on changes in nutrient supply and vice versa. It also showed that leaf cyanogenic glucoside content had been influenced differently by the combined influence of soil moisture content and nutrient supply. The leaf HCN levels obtained from the combined effects of soil moisture content and nutrient supply in *Salanga* and *Kiroba* are shown in Figs 1 and 2.

**Table 4.** F-test probability values for the two-way ANOVA on the effects of soil moisture content and nutrient supply on leaf HCN in *Salanga* and *Kiroba*.

| Factor | *Salanga*<br>p-value | *Kiroba*<br>p-value |
| --- | --- | --- |
| Soil moisture content (M) | < 0.001 *** | < 0.001 *** |
| Nutrient supply (N) | < 0.001 *** | < 0.001 *** |
| M×N | 0.002 ** | < 0.001 *** |

\*\*\* Significant at $p < 0.001$ , \*\* significant at $p < 0.01$ , \* significant at $p < 0.05$ and NS not significant ( $p > 0.05$ ).

**Fig 1.**
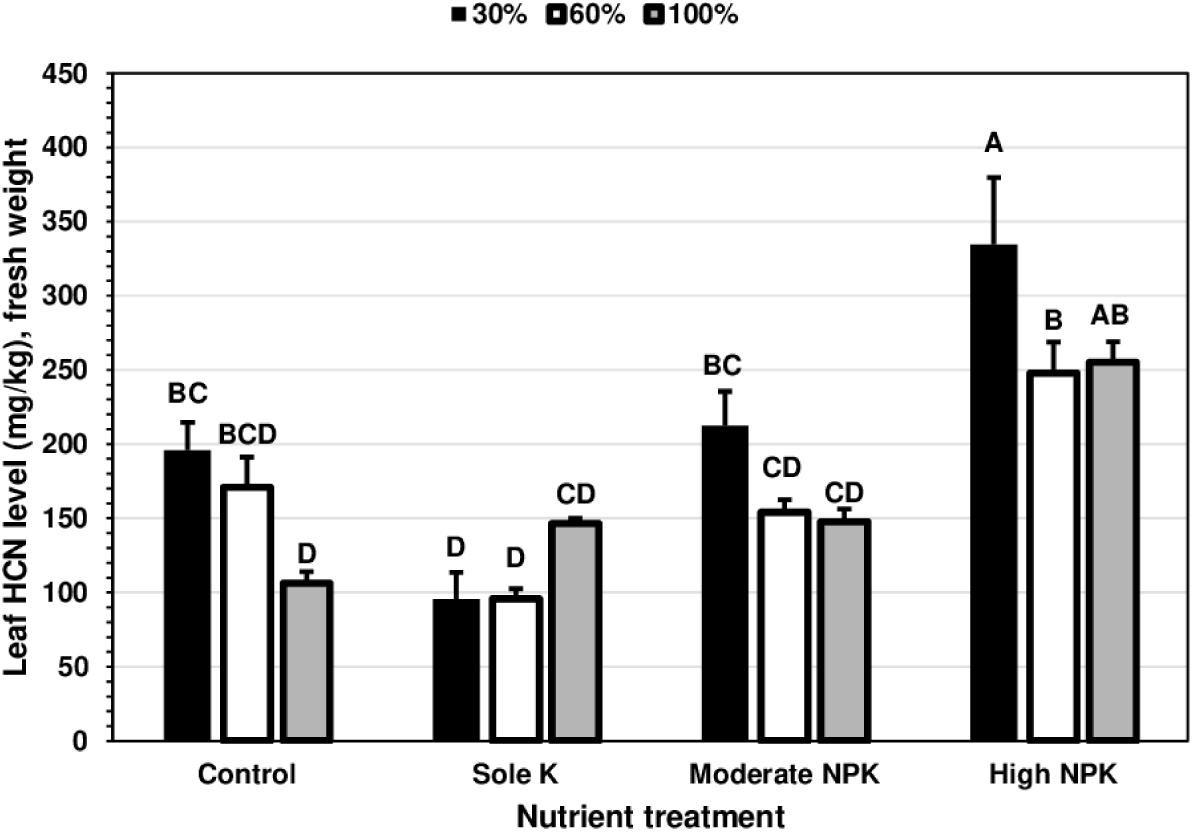
Leaf HCN in *Salanga* under the combined effects of soil moisture content and nutrient supply. For each variety, means (± SE) followed by even one same uppercase letter are not significantly different (Tukey’s test, p < 0.05). SE is the standard error of the mean.

**Fig 2.**
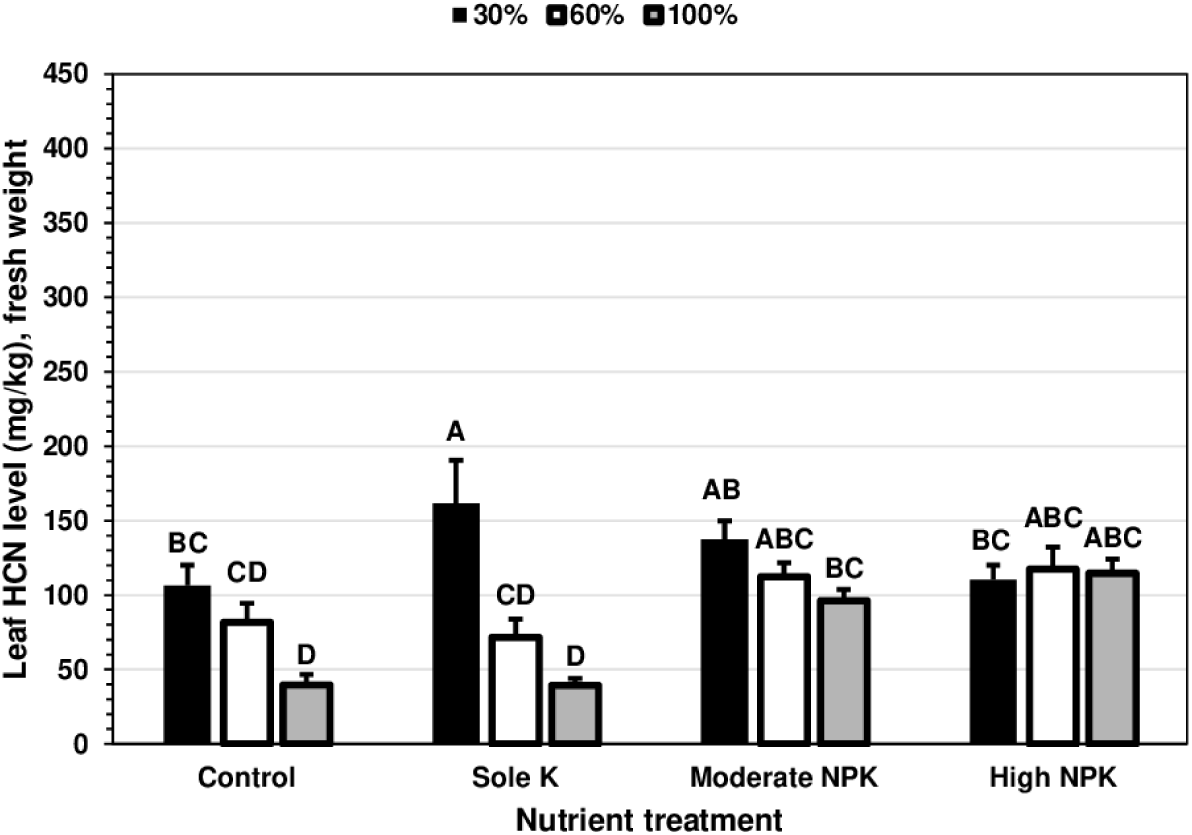
Leaf HCN in *Kiroba* under the combined effects of soil moisture content and nutrient supply. For each variety, means (± SE) followed by even one same uppercase letter are not significantly different (Tukey’s test, p < 0.05). SE is the standard error of the mean.

From Figs 1 and 2, it can be observed that changes in soil moisture could sometimes and sometimes not influence cyanogenic glucoside production in the two cassava varieties, all depending on soil nutrients conditions. When it did influence leaf cyanogenic glucoside contents, a significant increase in leaf HCN levels occurred as soil moisture conditions changed from well-watered to severe moisture stress. Severe moisture stress had increased leaf HCN levels of unfertilised *Salanga* by 1.8 times the levels attained under optimal soil moisture conditions. Due to severe moisture stress *Salanga* under the highest NPK treatment had its leaf HCN levels increased by 1.3 times the levels attained when it was moderately watered, but had similarly high leaf HCN levels as when it was well-watered. Furthermore, because of severe moisture stress, leaf HCN levels in *Kiroba* had increased by 2.7 times and 4.1 times the levels attained under optimal soil moisture conditions, when unfertilised and supplied with only K, respectively. Similar increments in cassava cyanogenic glucoside contents have been reported with increased soil moisture stress by other studies [39,40]. In one study, severely moisture stressed cassava plants had 2.9 times higher leaf HCN levels than levels found in leaves of well-watered plants [41].

Non-significant changes in leaf cyanogenic glucoside content were also just as common with changes in soil moisture under some soil nutrient conditions (Figs 1 and 2). When supplied with only K and with moderate levels of N, P and K, changes in soil moisture were observed to have no influence on the leaf cyanogenic glucoside content of *Salanga*. No influence on leaf cyanogenic glucoside contents were also observed in *Kiroba*, but unlike *Salanga*, this time it was with the supply of moderate and high amounts of N, P and K. One of the lowest leaf cyanogenic glucoside contents in *Salanga* were consistently obtained with sole K supply, no matter the soil moisture conditions. Sole K supply is commonly associated with reducing the cyanogenic glucoside content of cassava [19,42]. The non-significant influence of soil moisture content in *Salanga* under sole K supply, is thus beneficial as it would consistently result in low HCN levels in *Salanga*, regardless of changes in soil moisture conditions. However, unlike *Salanga*, the reducing effects of sole K supply in *Kiroba* were only evident under optimal soil moisture conditions and completely absent under severe moisture stress conditions. Well-watered conditions are thus needed for *Kiroba* to experience the reducing effects of sole K application. The results of the present study give a probable explanation for the inconsistent results obtained with sole K fertiliser application; that is, with it either reducing [19,42] or having no effect [13] on cyanogenic glucoside production in cassava. Increased cyanogenic glucosides in cassava with sole K application have however hardly been reported.

The non-significant response to changes in soil moisture in *Salanga* supplied with moderate amounts of N, P and K (Fig 3), is similarly as beneficial for reducing cyanogenic glucosides as sole K supply in this variety. This is because regardless of changes in soil moisture conditions, leaf HCN levels in *Salanga* supplied with moderate amounts of N, P and K had consistently remained at levels close to those attained with sole K supply. Reductions in cyanogenic glucosides with moderate applications of N, P and K have also been reported in other studies [2,12]. Moderate applications of NPK fertiliser (with N ≤ 50 kg/ha) could hence be useful for reducing cyanogenic glucoside levels in *Salanga* and in varieties like it, while high applications would only increase cyanogenic glucoside content to undesirable levels as was also seen in *Salanga* supplied with high rates of N, P and K, irrespective of soil moisture content. Some studies have however reported non-significant responses to N, P and K supply with moderate levels of N, P and K [43] and even with high levels of N, P and K [13], highlighting that there are further differences in how cassava varieties respond to soil nutrient supply. Reductions in leaf cyanogenic glucosides with the non-significant responses to changes in soil moisture by *Salanga* supplied with moderate amounts of N, P and K were however absent in *Kiroba*. This is because *Kiroba* had attained high leaf cyanogenic glucoside contents when supplied with both moderate and high amounts of N, P and K, close to the levels attained under severe moisture stress with sole K application. The increased supply of N, P and K even when supplied in moderate amounts would hence result in high cyanogenic glucoside levels in *Kiroba*, regardless of soil moisture conditions.

Taking a closer look at Table 4, it can also be seen that when moderately moisture stressed and when well-watered, leaf HCN levels of unfertilised *Salanga* were statistically just as low as levels attained under sole K supply. Thus, in the absence of severe water stress, not fertilising *Salanga* had similar reducing effects on leaf HCN like sole K supply. In the absence of severe water stress, not applying fertiliser was hence efficient at reducing leaf HCN levels of *Salanga*, but a contributor to increased cyanogenic glucosides in this variety under severe moisture stress. In the same way, not fertilising *Kiroba* would be just as beneficial at reducing cyanogenic glucosides in this variety in the absence of severe water stress. This is seen by the low leaf HCN levels attained by unfertilised *Kiroba* under well-watered conditions; its leaf HCN levels were significantly similar to the lowest leaf HCN levels it had attained when it was supplied with only K and had been kept well-watered. Leaf HCN levels in *Kiroba* however increased to levels similar to those attained with the moderate and high supply of N, P and K. The interaction effects of poor soil fertility and severely low soil moisture conditions could be contributing factors to the high cyanogenic glucoside levels observed during periods of water stress, in areas affected by cassava cyanide intoxication in sub-Saharan Africa. As previously mentioned, in addition to being prone to water stress, areas prone to cassava cyanide intoxication in sub-Saharan Africa are predominantly nutrient poor [5–7].

It is however important not to forget that the degree to which cyanogenic glucosides increase or decrease with changes in soil moisture content and nutrient supply, in both *Salanga* and *Kiroba*, is also dependent on their individual genetic control of cyanogenic glucoside production. The genetic control of cyanogenic glucoside expression in a variety, limits the possible range of cyanogenic glucoside contents attained by a variety as it is being influenced by its growing environment. According to how high and low their leaf cyanogenic glucosides had become under the same combined effects of soil moisture and nutrient supply, the sweet cassava variety *Kiroba* was seen to have a lower expression of cyanogenic glucosides (leaf HCN ranged from 39.3 mg/kg to 161.5 mg/kg) compared to the bitter cassava variety *Salanga* (leaf HCN ranged from 95.5 mg/kg to 334.5 mg/kg) (Fig 3).

The results of the present study demonstrate why responses are sometimes seen with NPK fertiliser application and why they are sometimes not observed. For instance, when *Kiroba* was kept well-watered or severely moisture stressed, significant responses were seen with the application of the NPK fertilisers. However, no responses were observed with NPK fertiliser application when *Kiroba* was kept moderately moisture stressed. On the other hand, *Salanga* always responded to NPK fertiliser application at all soil moisture levels.

### A holistic description of the cyanogenic characters of *Salanga* and *Kiroba*

Considering how soil moisture content and nutrient supply both influenced cyanogenic glucoside production in *Salanga*, the variety can be described as a cassava variety that would always produce high levels of cyanogenic glucosides on highly fertile soils. This is due to its sensitivity to a high supply of soil nutrients, irrespective of soil moisture conditions. *Salanga* is however a cassava variety that would do better on moderately fertile soils as it consistently acquires low cyanogenic glucosides when grown on such soils, regardless of soil moisture conditions. On the other hand, with regard to cyanogenic glucosides, *Kiroba* can be described as a cassava variety that is better adapted to nutrient poor soils and optimal soil moisture conditions. This is because cyanogenic glucoside levels in *Kiroba* are consistently increased by severe water stress, irrespective of soil nutrient conditions and also because cyanogenic glucosides in *Kiroba* get easily increased by even the slightest improvement of soil fertility.

## Conclusion

Using leaf HCN levels, the study managed to establish that nutrient supply had an equally important role in influencing cyanogenic glucoside production in cassava, just as varieties and soil moisture content. The study also revealed that cyanogenic glucoside production in cassava cannot be adequately described by the individual effects of varieties, soil moisture content and nutrient supply or by the interactive effects of varieties and soil moisture content alone. This is because cyanogenic glucoside production in various cassava varieties is differently influenced by the combined effects of soil moisture content and nutrient supply and also because the effects of either soil moisture content or soil nutrient supply on cyanogenic glucoside production in cassava varieties is dependent on the changes of the other factor.

The findings of the study highlight the need to pay greater attention to soil fertility and not only to differences in varieties and climatic conditions, when determining agronomic solutions for preventing the occurrence of high (toxic) levels of cyanogenic glucosides in cassava. This means that actual soil nutrient levels on fields need to be taken into consideration, in addition to defining soil fertility by soil classification names. The revealed information will be useful for breeding cassava varieties that will consistently give low root cyanogenic glucosides in any environment. Further research using root HCN levels, instead of leaf HCN levels is however needed. The findings of the present study additionally need to be verified under field conditions.

## Acknowledgements

Debts of gratitude firstly go to all the funders of the research, they made the work possible. Many thanks also go to the staff at the Root and Tuber Department at Naliendele Agricultural Research Institute (NARI), and the staff at the Department of Soils and Geological Sciences and also at the Horticulture Department at Sokoine University of Agriculture (SUA) for their technical support. We are additionally grateful to Dr R.H. Howeler and Dr J.H. Bradbury (Australian National University) for their useful insights on conducting cassava pot experiments and on carrying out cassava cyanide analysis, respectively. Lastly, special thanks go to Dr B.M. Humbel (University of Lausanne) for his encouragement throughout the entire study.

